# Visualizing the stochastic gating of Na^+^/H^+^ antiporters in single native bacteria

**DOI:** 10.1101/2022.07.03.498436

**Authors:** Yaohua Li, Haoran Li, Jia Gao, Ben Niu, Huan Wang, Wei Wang

## Abstract

Na^+^/H^+^ antiporters are the major secondary transporters that regulate pH and sodium homeostasis by enabling transmembrane exchanges of Na^+^ with H^+^ in opposite directions, both are essential cations. Although their crystal structures and functions have been well characterized^1-3^, the transport dynamics of Na^+^/H^+^ antiporters during action in living cells remained largely unexplored. Herein, intermittent blinking of the spontaneous bioluminescence from single native bioluminescent bacteria, *P. phosphoreum*, was reported, investigated and attributed to the stochastic gating of Na^+^/H^+^ antiporters between the active and inactive conformations. Each gating event caused the rapid depolarization and recovery of membrane potential within several seconds, accompanying with the intermittent bioluminescence blinking due to the transient inhibitions on the activity of respiratory chain. Temperature-dependent measurements further revealed that the conformational change was thermodynamically driven with an activation energy barrier of 20.3 kJ/mol. While the stochastic gating of ion channels has been well understood for decades^4,5^, this study uncovered the stochastic gating dynamics of Na^+^/H^+^ antiporters, another major pathway for ion transmembrane transports, in single native bacteria without any genetic engineering or chemical labeling. It also opened the door for *P. phosphoreum* to serve as new model bacteria for reporting the physiological and metabolic status with spontaneous bioluminescence emission.

## Introduction

Among many members in the sodium/proton antiporter family, NhaA is probably the best characterized one with available crystal structures and rational mechanisms of action^1,2^. As one kind of electrogenic antiporters, NhaA effluxes one Na^+^ against the concentration gradient while influx two H^+^ driven by the proton motive force. A major feature of NhaA (also shared by many other members^3^) is that its activity is highly regulated by cytoplasmic pH. Previous studies in reconstituted proteoliposomes^6^ and isolated membrane vesicles^7^ have revealed a dramatic increase of transport efficiency by ∼2000 times with a pH rise from 6.5 to 8.5. A cluster of residues forms a pH-sensitive domain whose conformational change upon deprotonation exposes the otherwise buried binding site of Na^+^, and thus activates the antiporter function^8^. While intensive experimental and theoretical efforts have been made to clarify the conformations at both inactive (low pH) and active (high pH) states, little is known regarding the dynamics of such conformational switching in living cells^9^. For instance, at single NhaA molecules level, it remains an open question whether the increase in pH results in a series of intermediate conformations with gradually enhanced but stable turn-over frequency, or it leads to the higher duty ratio of the active conformation during stochastic switching between two different conformations (*i*.*e*., a gating mechanism often observed in ion channels). In order to address this question, a capability for continuously measuring the activity of single NhaA molecules in living cells is highly desirable but it has yet to be demonstrated.

Single molecule dynamics have significantly advanced the fundamental understandings on the mechanisms of many important biomolecules such as ion channels^5,10^, enzymes^11^ and membrane receptors^12^, since they captured the dynamic processes of these molecules during actions, rather than the static pictures. Single channel patch clamp was probably the earliest technique that allowed for recording the activity of single molecules as a function of time, which revealed the discrete and stochastic gating of single ion channels between the open and closed states at millisecond timescales^4^. Although transporters serve as another major pathway for transmembrane exchange of ions, patch clamp is not sensitive enough for studying transporters because the transport efficiency (number of ions per molecule per second, *i*.*e*., current) of transporters is often 4-5 orders of magnitude lower than that of ion channels^7^. Single molecule Förster resonance energy transfer (smFRET) is so far the most frequently adopted technique to unravel the turn-over dynamics of many transporters (other than NhaA)^13-23^. When labeling a single transporter molecule with a donor and an acceptor at different sites, the conformational change altered the distance between the two fluorophores, which was revealed by the temporal fluctuations in FRET efficiency^24^. However, smFRET often required double labeling for signal transduction, which were not only labor intensive but also introduced inevitable disturbance to the structure and dynamics of the proteins of interest. It is particularly concerning for transporters whose functional sites are often buried within the transmembrane domains. As a result, while smFRET are powerful for *in vitro* studies of purified proteins, it remains rather challenging to investigate the dynamics of transporters in living cells.

Here we report the discovery of spontaneous and stochastic bioluminescence (BL) blinking of single native BL bacteria, *P. phosphoreum*. Stochastic gating of electrogenic NhaA antiporters caused the rapid depolarization and recovery of membrane potential within several seconds, which transiently suppressed the activity of respiratory chain, accompanying with the intermittent blinking of BL emission from single native BL bacteria, *P. phosphoreum*. While it has been known that weakly alkaline pH induces the conformational change of NhaA via the deprotonation of certain residues and activates its transport activity, visualization of the intermittent opening of NhaA unraveled that it was a thermodynamically-driven stochastic gating process with a subtle activation energy barrier of 20.3 kJ/mol under physiological conditions (native bacteria, pH 7.4). The improved transport efficiency at higher pH was achieved by increasing the duty ratio of active conformation. In addition to the well-known effects of NhaA on the homeostasis of H^+^ and Na^+^, this study brings an example of electrogenic NhaA to regulate membrane potential of single bacteria in spontaneous and stochastic manners. Moreover, because the signal transduction solely relied on a BL system that was endogenously expressed in native bacteria, this study also opens up a promising possibility for *P. phosphoreum* to serve as new model bacteria for rapidly and sensitively reporting the physiological and metabolic states (such as membrane potential) via spontaneous BL emission.

### BL blinking of single native bacteria

We first report on the phenomenon and features of BL blinking of single native bacteria *P. phosphoreum*. Bacteria were cultured according to conventional methods, harvested at the exponential growth stage, and incubated on a quartz coverslip that was previously functionalized with poly-D-lysine to enable the deposition and immobilization prior to experiments. A representative bright-field image of surface-attached sub-monolayer bacteria is displayed in Fig. 1a. *P. phosphoreum* is a neutrophilic Gram-negative bacteria that are able to spontaneously emit light as a result of the BL reactions involving the enzymatic oxidation of reduced flavin mononucleotide (FMNH_2_) and fatty aldehyde with oxygen^25-28^. They are widely used for environmental toxicity assessments by evaluating the inhibition effect of environmental water samples on the BL intensity. The strain was thus commercially available and used as received without any genetic treatments. Correlative BL image of the same region-of-interest (ROI) is shown in Fig. 1b, which was captured on a conventional wide-field fluorescence microscopy in the absence of excitation light and filter sets. A bluish green pseudo-color was applied to illustrate the characteristic emission wavelength of 478 nm (Fig. S1).

**Fig. 1.**
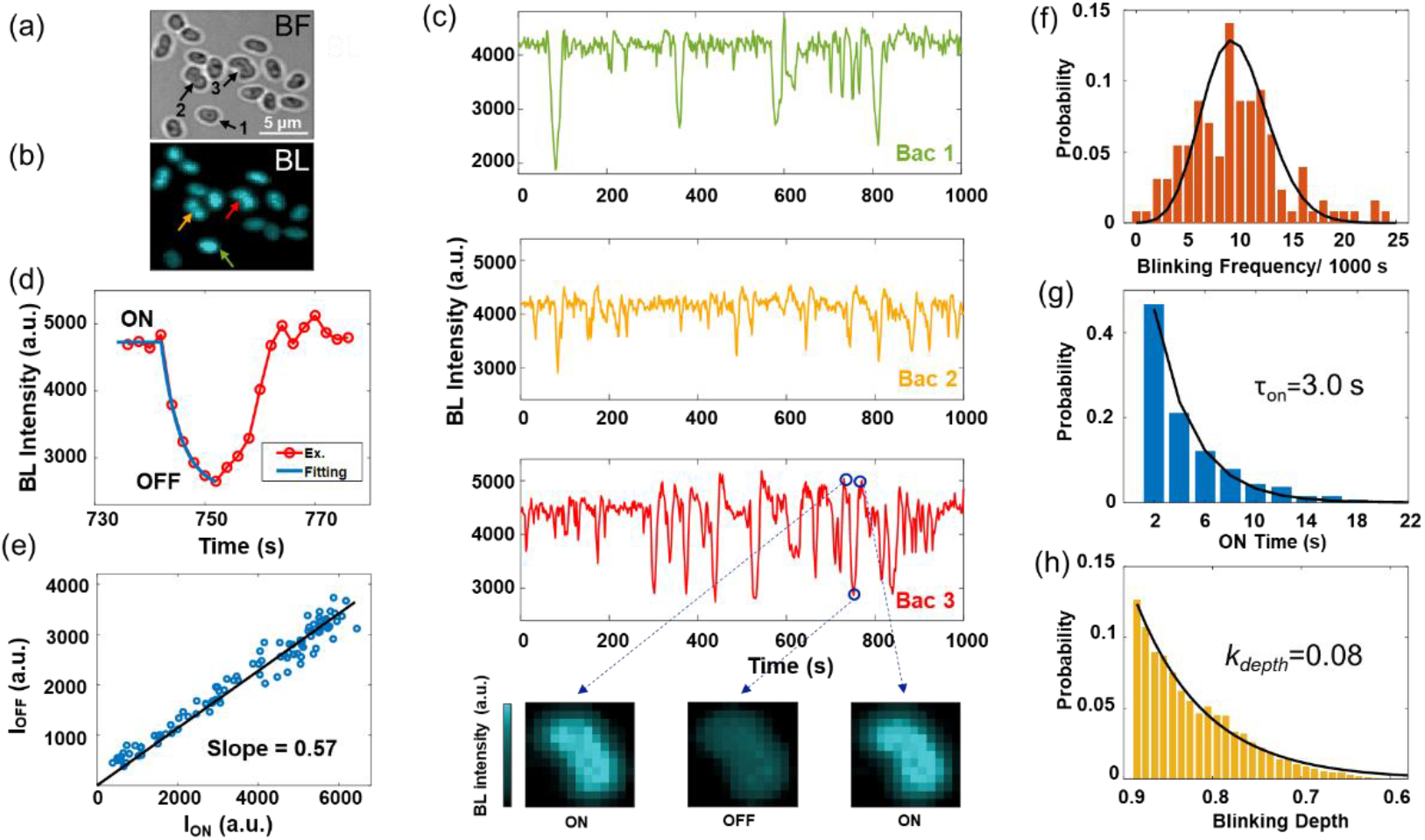
Phenomenon and dynamics of the intermittent BL blinking of single native bacteria. Correlative bright-field (a) and BL (b) images of surface-adhered *P. phosphoreum* (scale bar: 5 μm). (c) Representative BL curves of three individuals (marked in (a) and (b)). Zoom-in BL images of individual #3 during a blinking event are shown at the (c) bottom, together with the corresponding BL curve (d) and the inhibition ratio of each pixel (e). Histograms of the blinking event numbers (f), duration times of ON events (g), and blinking depths (h) of totally 1350 events observed in 126 bacteria in 1000 seconds.

When incubating the bacteria in a weakly alkaline medium (pH 8.0) and continuously recording the time-lapsed BL images, it was surprisingly found that almost all individual bacteria underwent stochastic and spontaneous blinking (Movie S1). Three representative BL intensity trajectories are provided in Fig. 1c, exhibiting several features in the blinking dynamics. (1) The blinking events of adjacent individuals were mostly independent. This feature excluded the dynamic fluctuation of local oxygen concentration (Fig. S2a). In some rare cases, two closely adjacent individuals exhibited synchronized trajectories (Fig. S2b), suggesting they were at the late stage of cell division and there were still efficient intercellular exchange of substances. (2) When zooming in a representative blinking event (Fig. 1d), BL intensity firstly exponentially decayed with a time constant of 3.6 seconds, then recovered within a comparable time scale. Multiple events in the same individual exhibited similar decay constants (Fig. S3). (3) For each blinking events, different subcellular locations/pixels exhibited similar intensity ratio (*I*_OFF_/*I*_ON_), *i*.*e*., blinking depth (Fig. 1e & Fig. S4), indicating a spatially uniform suppression of BL. (4) By applying a threshold of 0.9 on blinking depth to distinguish ON and OFF states in a trajectory (a popular method to quantitatively analyze blinking dynamics^29^), statistic analysis on 126 individuals exhibited, on average, 10 blinking events per bacterium in 1000 seconds (Fig. 1f). Large variations on the blinking frequency were observed among different individuals. The duration times of ON events (*τ*_ON_) followed an exponential distribution with a time constant of 3.0 seconds (Fig. 1g). The blinking depth also followed an exponential decay with a decay constant (*k*_*depth*_) of ∼0.08 (Fig. 1h). The frequent and rapid BL blinking within a few seconds (Fig. 1g), and the uniform blinking depth within the bacteria (Fig. 1c), indicated that it was caused by the dynamic changes in the substrate concentration, rather than the expression level of BL-related enzymes which was relatively slow and heterogeneous.

### Power spectral density analysis on blinking index

In order to extract an overall value, herein termed blinking index, to quantify the extent of BL blinking from a series of time-lapsed images, a power spectral density (PSD) method was introduced. This method has been developed to characterize the fluorescence blinking of single quantum dots^30^. Briefly, after obtaining the normalized BL curves of all single bacteria via automatic recognition and selection, averaged PSD was calculated to represent the distribution of signal fluctuation along frequency dimension (Fig. S5). According to the results in both spatial and temporal segmentations (Fig. S6), PSD analysis provided a reliable and quantitative manner to quantify the extent of blinking. A blinking index was further extracted by integrating the area within the frequency range of 0.01∼0.15 Hz in the averaged PSD curve.

### Dependence of BL blinking on pH and Na^+^

We subsequently conducted a series of experiments, including pH, cations, and specific inhibitors, to indicate that the stochastic gating of NhaA was responsible for the BL blinking. PSD curves under increasing pH from 5.0 to 9.0 were displayed in Fig. 2a. Plotting the blinking index as a function of pH revealed a sharp increase with increasing pH from 6.5 to 8.5 (Fig. 2b), which was well consistent with the previously reported influence of pH on the transport efficiency of NhaA^6,7^. Two movies captured at pH 6.5 and 8.5 were provided in the Supplementary Information for comparative illustration (Movie S2).

**Fig. 2.**
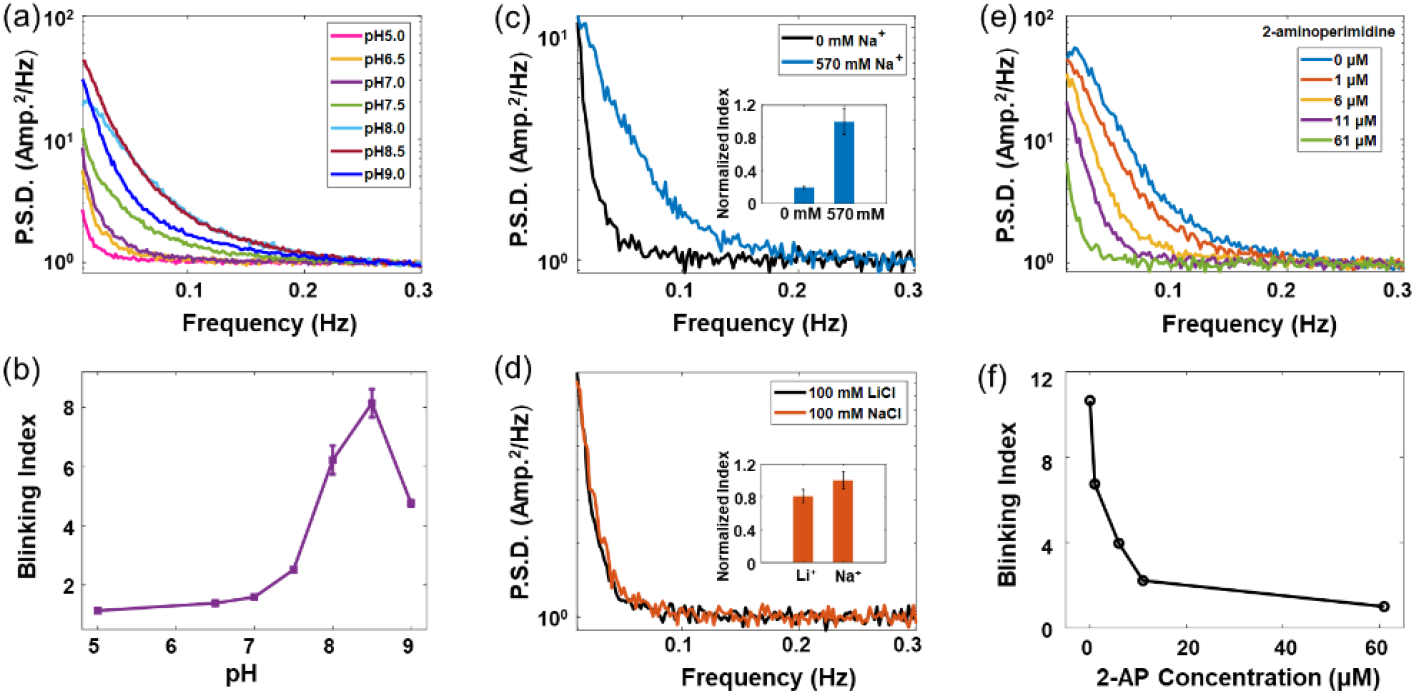
NhaA activity is associated with BL blinking. Influences of pH (a, b), Na^+^ (c), Li^+^ (d) and NhaA inhibitor, 2-AP, (e, f) on BL blinking, characterized by PSD curve (a, c, d, e) and blinking index (b, c inset, d inset, f), respectively.

Furthermore, a decrease in the blinking index by 80% was observed when reducing the Na^+^ concentration from 570 to 0 mM (Fig. 2c). The dependence of BL blinking on sufficiently high Na^+^ concentration was expected because the activity of NhaA relied on high intracellular Na^+^ concentration, which was regulated by extracellular Na^+^ concentration via Na^+^/solute symporter mechanism^31^. Furthermore, it has been reported that Li^+^ has an equivalent effect to Na^+^ on NhaA^3^. Indeed, replacement of Na^+^ with the same concentration of Li^+^ resulted in slightly decreasing blinking index (Fig. 2d). The significant dependence of BL blinking on both pH and Na^+^ concentration suggested the roles of NhaA. Genomic sequencing of as-used strain of *P. phosphoreum* also confirmed the existence of NhaA gene in its genome (Table S1).

### Effects of NhaA inhibitors

2-Aminoperimidine (2-AP) is a known inhibitor that specifically targets bacteria NhaA^32^. Fig. 2e and 2f show the PSD curves and the reducing blinking indices, respectively, with the increasing concentration of 2-AP. The half inhibition concentration (IC_50_) was 1 μM, which was consistent with literature reports on the inhibition concentration of 2-AP on NhaA. 5-(N, N-dimethyl)-amiloride (EIPA) was another popular inhibitors for Na^+^/H^+^ exchanger in mammalian cells. It is also reported to inhibit the activity of NhaA in marine bacterium *Vibrio parahaemolyticus*^33^. Our results indicated that EIPA suppressed the blinking index with similar IC_50_(6.5 μM) with literature reports (Fig. S7).

### Synchronized fluctuations in membrane potential

While the action of electrogenic NhaA might have altered both the intracellular pH (pH_i_) and membrane potential by exchanging one Na^+^ (efflux) with two H^+^ (influx), we clarified that the depolarized membrane potential, rather than the decreased pH_i_, was responsible for the BL blinking. As shown in Fig. 3a, when labeling the bacteria with tetramethylrhodamine methyl ester (TMRM), a widely-used FL probe for bacteria membrane potential^34^, correlative FL and BL trajectories of a single bacterium clearly showed synchronized drops in the blinking events. Such comparison not only demonstrated the BL blinking was accompanied with the transient depolarization of membrane potential, but also suggested BL could be an alternative and even better signal transduction format to readout the transient and weak fluctuations in membrane potential with a faster response (thus better temporal resolution) and higher sensitivity (Fig. S8). The effect of valinomycin (a K^+^-specific ionophore^3^) and carbonyl cyanide m-chlorophenyl hydrazine (CCCP, a proton-specific ionophore^34^) on BL blinking were further examined, which were popularly used to eliminate the transmembrane potential by collapsing the K^+^ concentration gradient and proton motive force, respectively. Both valinomycin (Fig. 3b) and CCCP (Fig. 3c) were found to efficiently reduce the blinking index, validating the roles of membrane potentials in BL blinking. Given a typical cell membrane capacitance density of 1 μF/cm^2^, surface area of 2 μm^2^, and a turn-over frequency of single NhaA of 1500 per second^3^, a net influx of 1500 positive charges in one second should reasonably depolarize the membrane potential by 10 mV. A typical ON event last for several seconds (Fig. 1g), corresponding to a depolarization by several tens of mV.

**Fig. 3.**
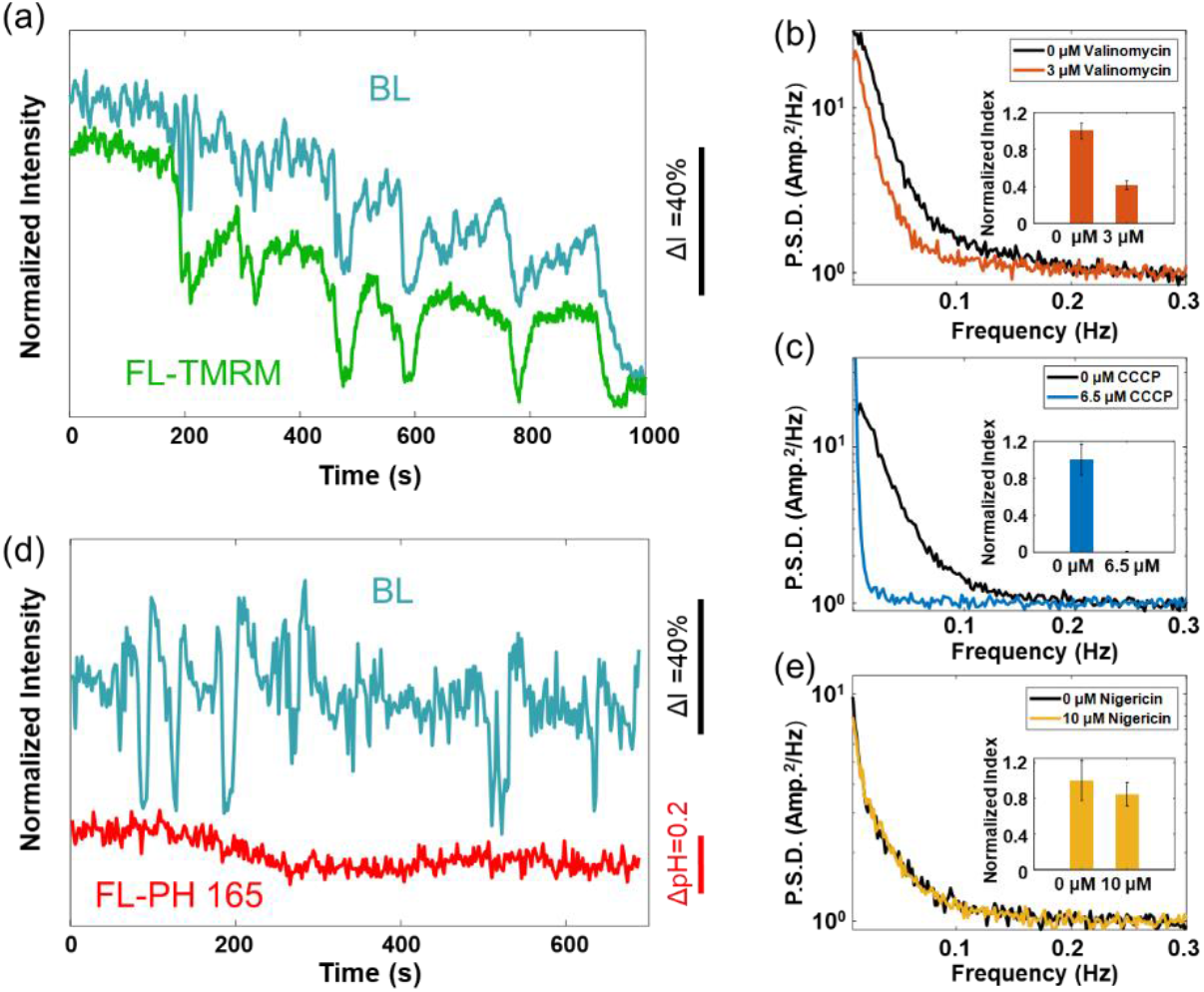
Transiently depolarized membrane potential is responsible for BL blinking. Correlative BL (blue) and FL (a: green, d: red) recordings when labeling the BL bacteria with FL probes for membrane potential (a, TMRM) and for intracellular pH (d, RatioWorks™ PH165 NHS ester), respectively. The influences of valinomycin (b), CCCP (c), and nigericin (e) on BL blinking are characterized by PSD curves and blinking index (insets), respectively.

### Intracellular pH remains stable during blinking

A ratiometric pH-sensitive fluorophore, RatioWorks™ PH165 NHS ester, was further introduced to investigate dynamics of pH_i_ during blinking events. As shown in Fig. 3d, while the BL trajectory exhibits frequent and obvious blinking features, the correlatively recorded pH_i_ remains stable. This point was further supported by the comparable blinking index between the absence and presence of 10 μM nigericin (Fig. 3e), a proton ionophore that preferably eliminated the transmembrane ΔpH while maintaining the membrane potential^35^. We clarify that the probe was indeed capable for reporting the pH_i_, because in some rare individuals (less than 3% of the population), synchronized drops in pH_i_ trajectory with rather small amplitudes were indeed observed at the moment of BL blinking (Fig. S9). The fact that pH_i_ remained constant during the BL blinking demonstrated that, the cytoplasm owned a greater buffering capacity to keep the pH_i_ stable. Buffering capacity of neutrophilic bacteria was previously reported to be 10∼100 nmol protons per pH unit per mg cell protein under physiological conditions^36^, thus requiring an influx of 10^5∼6^ protons to alter 1 pH unit (Fig. S9). Given the maximum transport efficiency of 1500 protons per second^3^, it was reasonable to understand that the activation of NhaA was insufficient to obviously alter the pH_i_.

### Activation energy barrier for conformational change

The spontaneous conformational change was driven by thermal fluctuation. When gradually increasing the temperature from 28 to 44 °C and plotting the histograms of blinking depths, it was found that the probability with larger inhibition ratio (thus smaller blinking depth) was increased (Fig. 4a and Fig. S10), indicating the higher duty ratio of active state and thus the larger rate of conformational change from the inactive to active state. According to Arrhenius’ equation, the temperature-dependent gating dynamics thus allowed to estimate the activation energy barrier (*E*_a_). Because the blinking depth was determined by *τ*_on_ (longer *τ*_on_ led to larger blinking depth, Fig. S11), plotting the logarithm of *k*_depth_ as a function of 1/T resulted in an *E*_a_ of 20.3 kJ/mol (Fig. 4b).

**Fig. 4.**
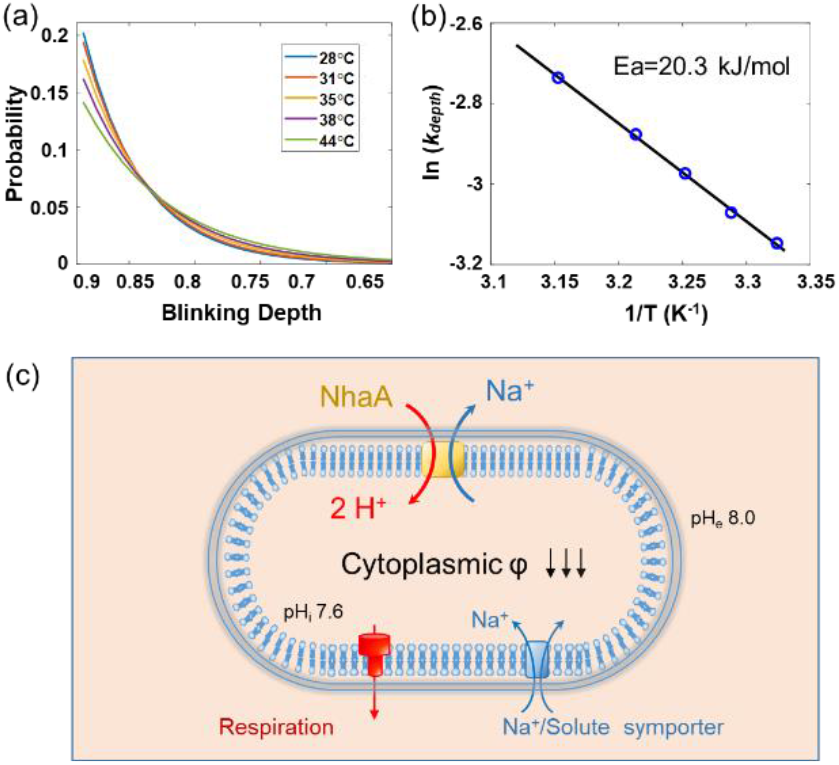
Activation energy barrier of NhaA conformational change. (a) Dependence of the blinking depth on temperature, T. (b) Activation energy of the NhaA conformational change is determined by plotting ln(*k*_*depth*_) as a function of 1/T. (c) Schematic illustration on the mechanism of BL blinking.

## Discussion

A mechanism was therefore reached by putting all the pieces together (Fig. 4c). Since an individual *P. phosphoreum* has a surface area of ∼2 μm^2^, it is possible that there are only one or a few NhaA molecules expressed in the membrane to induce discrete blinking events, rather than continuous BL fluctuations^37^. This point was further supported by the observation that different blinking events exhibited similar decay constants in the trajectory curves (Fig. S3). The stochastic switching from the inactive to active state enabled the net influx of, in maximum, 1500 positive charges per second, resulting in the depolarization of membrane potential by several tens of mV depending on the duration time of ON event. Because the proton transport was in fact driven by the proton motive force, the depolarized membrane potential should reduce the transport efficiency until a new equilibrium was established. Such negative feedback was therefore responsible for the exponential decay in the membrane potential and thus BL intensity (Fig. 1d). The variation in the inhibition depths in different blinking events (Fig. 1h) was caused by the different inhibition times that were not long enough to reach the equilibrium. Therefore, the histogram of blinking depth was directly correlated with the ON duration time (Fig. S11). Moreover, rise in cytoplasmic pH_i_ (in accordance with the extracellular medium pH_e_, Fig. S12) led to significantly increased duty ratio of the active state, as evidenced by the clear dependence of blinking index on pH (Fig. 2b). The active state would not last forever. Instead, a spontaneous switching from the active to inactive state disabled the influx of H^+^ and stopped the depolarization. The rapid recovery of membrane potential and BL intensity indicated an effective mechanism to pump out the net positive charge, likely by H^+^-ATPase. In order to verify this point, two H^+^-ATPase inhibitors, oligomycin, and N, N’-dicyclohexylcarbodiimide (DCCD), were examined (Fig. S13). It was found that the presence of oligomycin and DCCD efficiently inhibited the blinking index by 40% and 55%, with the corresponding inhibition concentrations of 20 μM and 1 mM, respectively.

The dependence of BL intensity on membrane potential was attributed to the regulation of respiratory chain activity. The depolarized membrane potential suppressed the activity of FMN reductase and decreased the production rate of FMNH_2_, which was a major substrate of luciferase. BL emission resulted from a cascade reaction involving multiple enzymes including flavin reductase, fatty acid reductase complex, and luciferase^26^. The inhibition of flavin reductase immediately decreased the concentration of intermediate FMNH_2_ and thus the BL intensity. In order to examine the generality of such a mechanism, another BL bacteria, *Vibrio fischeri*, was investigated. Similar BL blinking was also observed in wild type *Vibrio fischeri*, which also exhibited a significantly enhanced blinking index with increasing pH from 7.0 to 8.0 (Fig. S14 & Movie S3).

## Acknowledgments

We thank the National Natural Science Foundation of China (Grants 21925403, 21874070), and the Excellent Research Program of Nanjing University (Grant ZYJH004) for financial support.

## Author contributions

Y.H.L. and W.W. designed the project and wrote the paper. Y.H.L. carried out most of the BL experiments and analyzed the data. H.R.L. developed the PSD analysis method. J.G. and B.N. helped the correlative BL and FL recordings. H.W. analyzed and disscussed the results and helped the treatments on bacteria. W. W conceived and supervised the project.

