## Supplement Fig.S1-14 for "Visualizing the stochastic gating of Na^+^/H^+^ antiporters in single native bacteria"

This PDF file includes:

Materials and Methods

Supplementary Figures S1 to S14

Supplementary Table S1

References

Other Supplementary Information for this manuscript includes the following:

Movie S1

Movie S2

Movie S3

### Table of Contents


#### 1. Materials and Methods

*1.1 Chemicals*
*1.2 Strains and culture conditions*
*1.3 Bacteria immobilization*
*1.4 Culture medium in Na+ and Li+ concentration experiments*
*1.5 Introduction of TMRM and RatioWorks™ PH165, NHS ester to cells*
*1.6 Temperature control*
*1.7 Microscope apparatus*

#### 2. Supplementary Figures

*Figure S1. BL emission spectrum*
*Figure S2. BL trajectories of adjacent individuals*
*Figure S3. Zoom-in trajectory of multiple blinking events*
*Figure S4. Consistent inhibition ratio of different pixels within single bacteria*
*Figure S5. Procedures for power spectral density analysis*
*Figure S6. Validations on the reliability of power spectral density analysis*
*Figure S7. Effects of NhaA inhibitors*
*Figure S8. Synchronized BL blinking and fluctuations in membrane potential*
*Figure S9. Occasionally synchronized BL blinking and fluctuations in intracellular pH*
*Figure S10. Distribution histograms of blinking depths at five temperatures*
*Figure S11. The dependence of* $\tau_{on}$ *and* $k_{depth}$
*Figure S12. Measurements of intracellular pH*
*Figure S13. Effects of* $H^+$*-ATPase inhibitors*
*Figure S14. BL blinking in wild type V. fischeri*

#### 3. Supplementary Movies

*Movie S1. BL blinking of P. phosphoreum*
*Movie S2. BL blinking in the solution of pH 6.5 and pH 8.5*
*Movie S3. BL blinking of V. fischeri*

#### 4. Supplementary Table

*Table S1. Gene and amino acid sequences of NhaA in P. phosphoreum*

#### 5. References

#### Materials and Methods

##### 1.1 Chemicals

NaCl, $Na_2HPO_4$, $KH_2PO_4$, and glycerol were obtained from Nanjing Chemical Reagent CO., Ltd. Yeast extract was obtained from Adamas Reagent, Ltd. Peptone was purchased from Qingdao Hope Bio-Technology Co., Ltd. Poly-D-Lysine was purchased from Sigma-Aldrich. Tetramethylrhodamine methyl ester (TMRM) and RatioWorks™ PH165, NHS easter were purchased from AAT Bioquest. 2-Aminoperimidine, 5-(N, N-dimethyl)-amiloride (EIPA), cariporide, carbonyl cyanide m-chlorophenyl hydrazine (CCCP), valinomycin, and nigericin were obtained from Shanghai yuanye Bio-Technology Co., Ltd. N, N'-dicyclohexylcarbodiimide (DCCD) was purchased from Shanghai Macklin Biochemical Co., Ltd. Oligomycin was purchased from Aladdin Holdings Group Co., Ltd.

##### 1.2 Strains and culture conditions

Wild type *P. phosphoreum T3* and *V. fisher* strains were used in this study, which were purchased from Institute of Soil Science, Chinese Academy of Sciences, and Beijing aiqing Technology Co., Ltd (No. AZF686300), respectively. Gene sequencing of *P. phosphoreum T3* was performed by Sangon Biotech Co., Ltd. (Shanghai, China) with Illumina HiSeq and PacBio RSII systems. Sequence assembly was achieved by SPAdes. PrInSeS-G is used for sequence correction. Gene functional annotation was based on CDD, KOG, COG, NR, NT, PFAM, Swissprot, and TrEMBL databases.

*P. phosphoreum T3* and *V. fisher strains* were cultured in the medium which contained 30.0 g/L NaCl, 5.0 g/L $Na_2HPO_4$, 1.0 g/L $KH_2PO_4$, 5.0 g/L yeast extract, 5.0 g/L peptone, and 3.0 mL glycerin with a pH of 7.0 ± 0.2. They were all cultured at 20 °C and shaken overnight at the speed of 200 rpm.

##### 1.3 Bacteria immobilization

The glass coverslip was pretreated with poly-D-Lysine (0.02 mg/mL) solution for 30 minutes and then washed with sterile water three times. After the glass coverslip was thoroughly air-dried, 100 μL bacterial solution was added to the chamber (made with polydimethylsiloxane) located on this glass coverslip. The bacterial solution was removed after 30 minutes and then gently rinsed with the culture medium to take away un-adhered bacteria in the suspension. Finally, the culture medium without nitrogen source (30.0 g/L NaCl, 5.0 g/L $Na_2HPO_4$, 1.0 g/L $KH_2PO_4$, 3.0 mL glycerin) was used to investigate bacterial blinking. The culture medium without nitrogen source can inhibit bacterial division and maintain stable BL for 1~2 hours.

##### 1.4 Culture medium for $Na^+$ and $Li^+$ concentration experiments

To completely remove $Na^+$ (0 mM) in the culture medium, NaCl was replaced for 500 mM glucose to partially compensate the osmotic pressure. The rest components contained 5.0 g/L $K_2HPO_4$, 1.0 g/L $KH_2PO_4$, 3.0 mL/L glycerin in the culture medium. The culture medium of 570 mM $Na^+$ contained 500 mM NaCl, 5.0 g/L $Na_2HPO_4$, 1.0 g/L $KH_2PO_4$, 3.0 mL/L glycerin. pH of these two medium solutions was adjusted to 8.0.

In order to examine the influence of $Li^{+}$, phosphate buffer was replaced for 50 mM Tris-HCl buffer solution to avoid precipitation formation (for example, $Li_3PO_4$). Other components were 100 mM LiCl, 400 mM glucose, 0.5 g/L KCl, 3.0 mL/L glycerin. The corresponding culture medium of $Na^{+}$ contained 100 mM NaCl, 50 mM Tris-HCl, 400 mM glucose, 0.5 g/L KCl, 3.0 mL/L glycerin. pH of these two medium solutions was 8.5.

##### 1.5 Introduction of TMRM and RatioWorks™ PH165, NHS ester to cells

In order to make a commercially-available membrane potential-sensitive fluorescent dye, TMRM, permeate into the bacterial cell wall, the bacterial suspension was pretreated with 10 mM chelating agent EDTA solution for 15 minutes. Then it was centrifuged to remove EDTA at the speed of 6500 rpm for one minute. These treated bacteria were incubated with 10 μM TMRM in PBS solution (containing 3% NaCl) for 1 hour. Finally, these bacteria were centrifuged and washed with the culture medium three times prior to immobilization.

RatioWorks™ PH165, NHS ester is a fluorescent dye that is suitable for ratiometric determination on intracellular pH. Bacteria were incubated with 10 μM PH165, NHS ester, in PBS solution (containing 3% NaCl) for 45 minutes. Then bacteria were centrifuged and washed with the culture medium three times prior to immobilization.

##### 1.6 Temperature control

We adopted the conductive indium tin oxide-coated coverslip (ITO, 15-30 Ω) as a resistor to heat the local medium surrounding the bacteria. The heating apparatus and calibration methods were slightly modified from our previous study[1]. According to Joule's law, applying different electric power can reach different local temperatures. The calibration curve between electric power and the local temperature was determined by two compounds, N, N'-dicyclohexylcarbimide ($C_{13}H_{22}N_2$), and lauric acid ($C_{12}H_{24}O_2$), with known melting points of 34 and 44°C, respectively. The melting process was observed under bright field imaging.

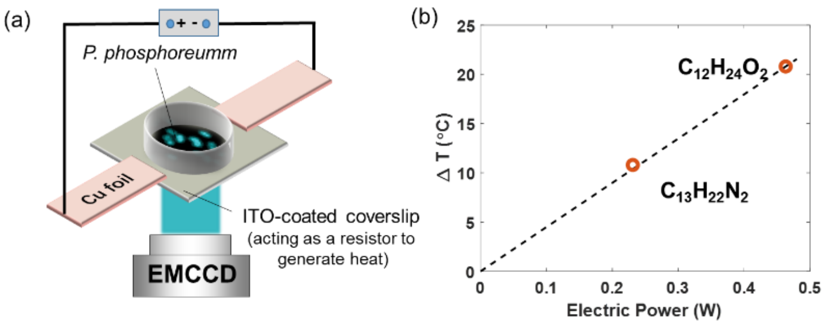


##### 1.7 Microscope apparatus

Bioluminescence of single bacteria was imaged with an Olympus IX83 inverted fluorescence microscope with a 60x oil immersion objective (N.A. =1.49) in the darkroom. All the excitation and emission filters and dichroic mirrors were removed from the light path, leaving a completely empty cube. The Olympus IX83-ZDC module can stabilize vertical focus. The emitted photons were collected by an electron multiplying charge-coupled device (EMCCD, Andor IXon Ultra 897, 512 × 512 pixels, 16 μm per pixel) camera. The exposure time (2 s for Fig. 1, and 1 s for Fig. 2-4), and an electron magnification (EM) gain value of 300 were used for recording BL images. Bright field imaging was illuminated with a standard halogen lamp.

Wide field fluorescence of the membrane potential dye, TMRM, was excited with 532-nm laser (Coherent OBIS, LS-532). The power of the continuous-wave (CW) 532-nm laser was 10 μW. Excitation filter 532/18 nm, emission filter long-pass 550 nm, and dichroic mirror 550 nm (Semrock) were utilized for capturing the fluorescence images. 488-nm laser (Coherent OBIS, LX-488) was used for exciting the fluorescence of ratiometric pH-sensitive dye, RatioWorks™ PH165 NHS ester, with a power of 10 μW. Excitation filter 481/18 nm, emission filter long-pass 650 nm, and dichroic mirror 550 nm (Semrock) were utilized for capturing the fluorescence images. The camera parameters of these two FL dyes were the same as BL imaging (exposure time: 1 s; EM gain: 300).

Correlative recording of BL and FL signals was achieved by automatically switching the microscope filter cube between a BL cube (empty) and a FL cube (with corresponding filter sets), and correspondingly switching the laser OFF/ON. This programmable process was operated by the Cellsens software (Olympus) to control all hardwares integrated into a real-time controller apparatus (RTC, Olympus). All BL and FL images were analyzed with ImageJ software and home-written MATLAB codes.

### Supplementary Figures

#### 2.1 BL emission spectrum

The suspension of *P. phosphoreum* (Fig. S1a) was diluted to $OD_{600\ nm}$ (optical density) value of 0.2. Emission spectrum of the diluted bacterial suspension was measured at room temperature with a fluorescence spectrophotometer under luminescence mode (F-7000, Hitachi). Representative BL emission spectrum was shown in Fig. S1b below. Maximal emission wavelength of *P. phosphoreum* was 478 nm.

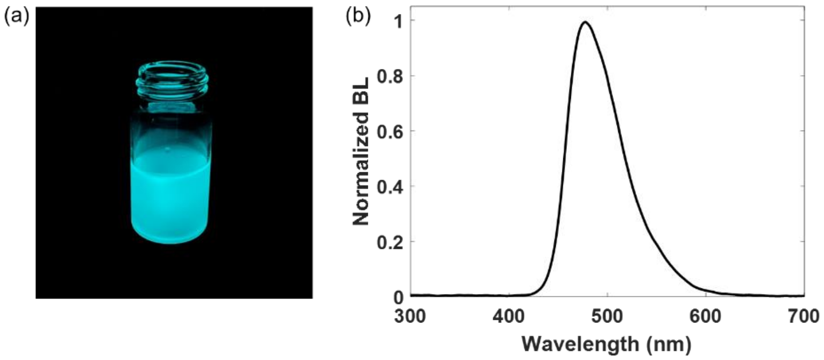

Fig. S1. (a) True-color photograph of the bacterial suspension. (b) The BL emission spectrum of *P. phosphoreum*.

#### 2.2 BL trajectories of adjacent individuals

For the majority of the adjacent bacteria, their BL trajectories were found to be independent from each other (Fig. S2a). This feature excluded the dynamic fluctuation of local oxygen concentration. The large diffusion coefficient of oxygen ensured that it only takes less than 1 ms for oxygen molecules to diffuse by 1 μm. Two individuals apart by 1 μm should experience nearly identical local concentration of oxygen at a timescale of 1 second.

However, in some rare cases, two adjacent individuals could exhibit highly synchronized blinking trajectories, as shown in Fig. S2b. Under such scenarios, the two individuals were always under close contact, suggesting they were at the late stage of cell division and there remained an efficient exchange of substances between them.

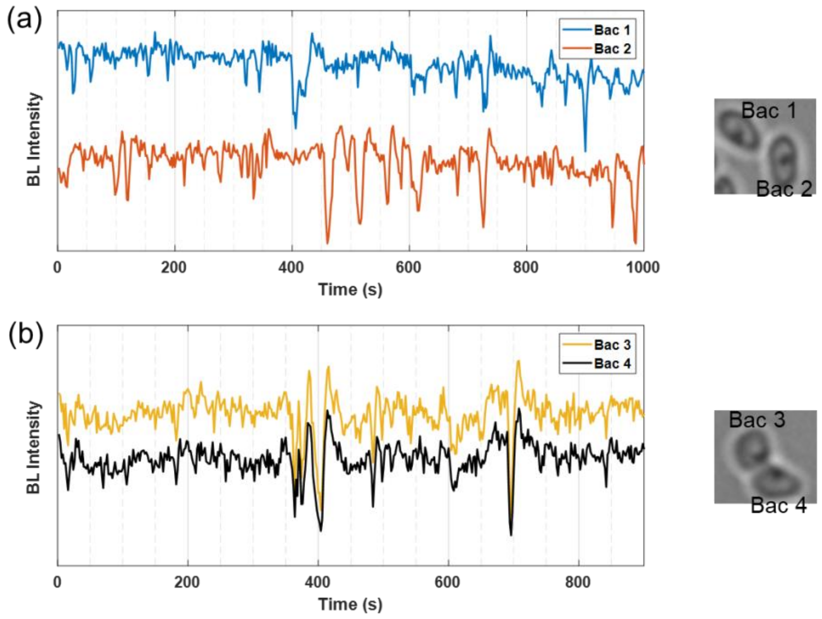


Fig. S2. (a) Blinking trajectory of a single bacterium was usually independent of that of its neighbors. (b) Synchronized trajectories were occasionally observed. Corresponding bright-field images are shown on the right panel.

#### 2.3 Zoom-in trajectory of multiple blinking events.

When zooming in three BL blinking events in Fig. 1c (Bac #3), it was found that each consisted of an exponential decrease and a subsequent recovery within similar time scales (Fig. S3). Comparable decay constants were obtained among different blinking events. It supported the hypothesis that different blinking depths resulted from the different duration times ($\tau_{on}$) – longer $\tau_{on}$ led to larger blinking depth.

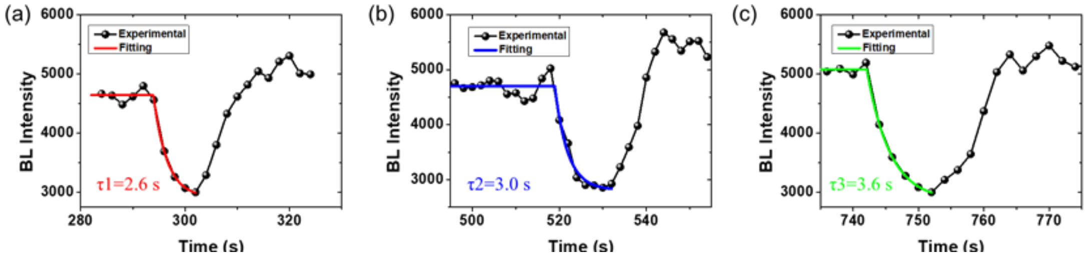


Fig. S3. Zoom-in BL trajectories of three blinking events recorded on a single bacterium.

#### 2.4 Consistent inhibition ratio of different pixels within single bacteria

Fig. S4a shows a representative BL trajectory consisting of several blinking events. Six snapshots of BL images are displayed below the curve, including four images in ON states and two images in OFF states during two blinking events. For the first event occurring between 56$^{th}$ and 84$^{th}$ seconds, a linear dependence of OFF state intensity (y-axis) with ON state intensity (x-axis) is observed for each pixel (Fig. S4b), exhibiting a slope of 0.74. This value was defined as the blinking depth. It meant that the BL intensity was uniformly inhibited by 26% regardless of the subcellular location. For the second event, uniform inhibition is also observed with a larger blinking depth of 0.66 (Fig. S4c). These results indicated that, while the blinking depth was spatially uniform within a single bacterium during one blinking event, different blinking depths were observed in different blinking events. It was likely due to the different duration times ($\tau_{on}$) – longer $\tau_{on}$ led to larger blinking depth.

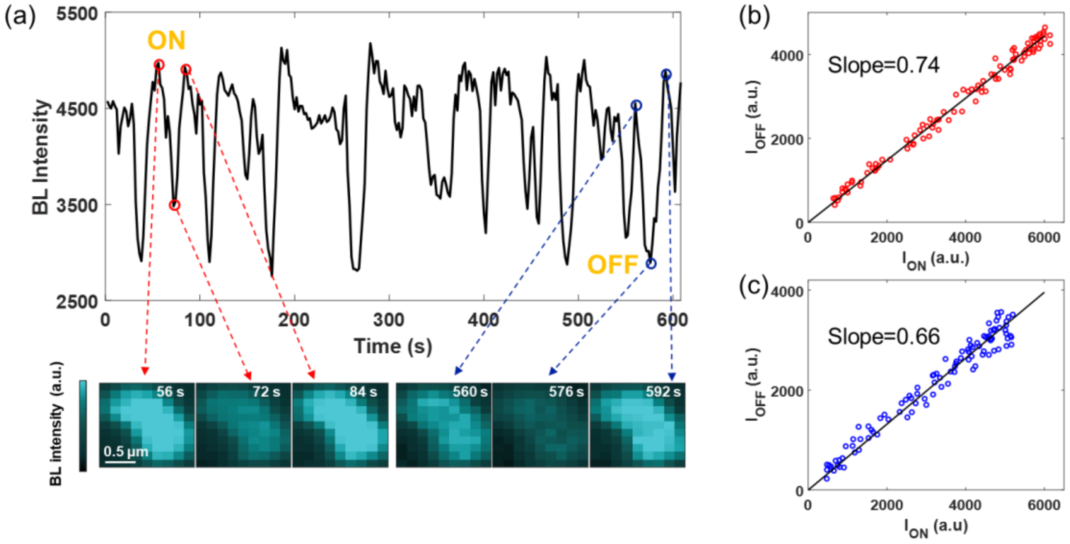


Fig. S4. (a) BL intensity curve and the corresponding BL image snapshots of a single bacterium. (b)-(c) Linear dependence of the BL intensities of each pixel in ON state with those in OFF states in blinking events occurring at the 72$^{th}$ second (b, red dots) and the 576$^{th}$ second (c, blue dots).

##### 2.5 Procedures for power spectral density analysis

In order to quantify the extent of BL blinking, a power spectral density (PSD) method was introduced, consisting of four steps. First, an averaged BL image was obtained from a series of time-lapsed BL images (for instance, 600 images in 10 minutes). Second, single bacteria were automatically identified and located by applying the classical object recognition algorithm to the averaged BL image (with enhanced signal-to-background ratio). Third, BL intensity curves of each single bacteria were extracted as shown in Fig. S5c. Finally, power spectral density curves of each BL trajectories were calculated and overlaid in Fig. S5d (gray background). The averaged PSD curve was shown in Fig. S5d (red curve). The averaged PSD curve was integrated within the frequency range from 0.01 to 0.15 Hz, leading to a blinking index value to quantify and compare the extent of blinking between different experiments.

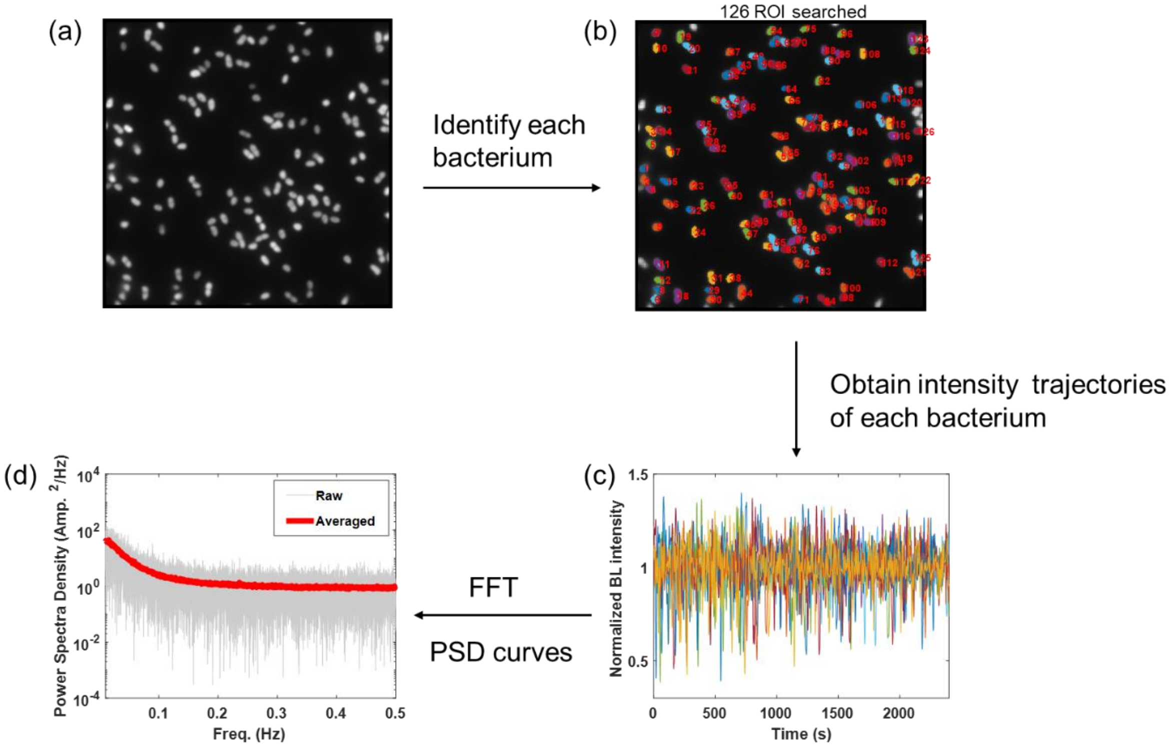


Fig. S5. (a) The averaged BL image. (b) Each single bacterium was identified, located and marked by corresponding color and numbers. (c) BL trajectories of all 126 bacteria were normalized and overlaid. (d) The corresponding 126 PSD curves are overlaid (gray background) and averaged (red curve).

##### 2.6 Validations on the reliability of power spectral density analysis

Two tests were examined to evaluate whether the above-mentioned PSD method (Fig. S5) was able to extract a reliable blinking index or not. The basic criterium was that the same blinking index values should be obtained from the same experiment regardless of the selection of ROIs. First, as shown in Fig. S6a, the original BL images were divided into 9 ROIs. PSD analysis was subsequently applied to each ROIs. It was found that the nine PSD curves were almost coincident and the nine blinking indices were consistent with each other, exhibiting a relative standard deviation of 8%. Besides spatial segmentation, temporal segmentation was also examined. As shown in Fig. S6b, a 45-min movie was temporally divided into 9 periods (5 minutes each). PSD analysis was applied to each period. It was similarly found that the nine PSD curves were comparable with each other. The nine blinking indices were consistent, exhibiting a relative standard deviation of 7%.

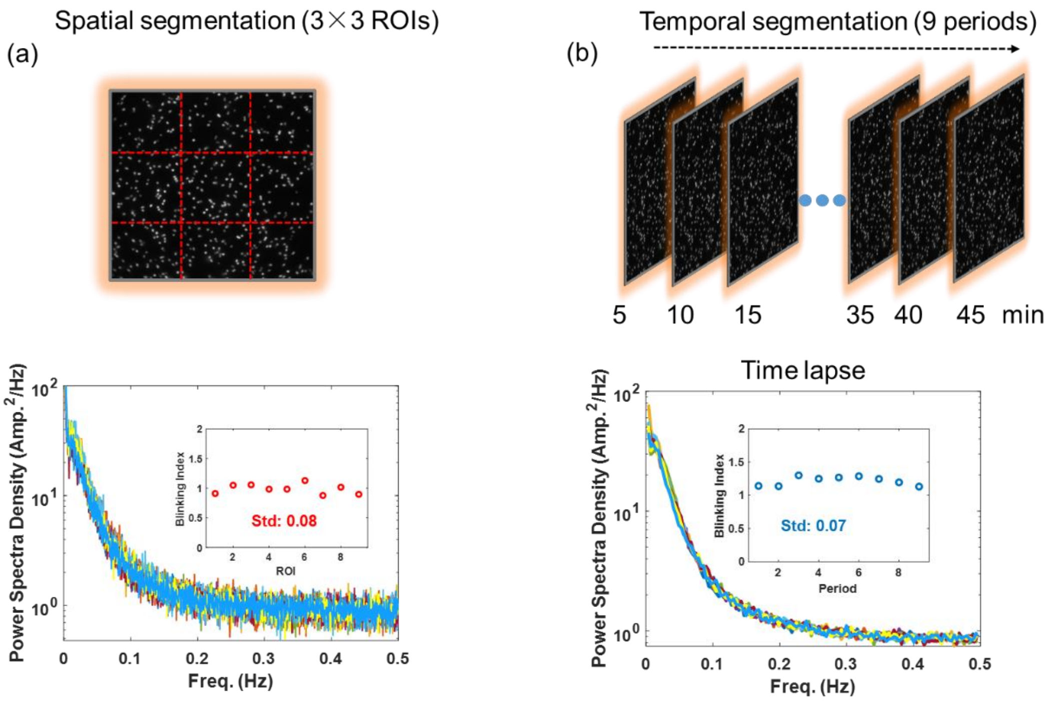


Fig. S6. PSD curves and the corresponding blinking indices of nine ROIs via spatial (a) or temporal (b) segmentations.

#### 2.7 Effects of NhaA inhibitors

5-(N, N-dimethyl)-amiloride (EIPA) is one kind of inhibitor for $Na^+/H^+$ exchanger in mammalian cells. It was also reported to inhibit $Na^+/H^+$ antiporter in bacterial cells. It was found that the presence of EIPA obviously reduced the blinking index with the increasing concentration of inhibitors (Fig. S7). The half-maximal inhibitory concentration, $IC_{50}$, of EIPA was 6.5 μM.

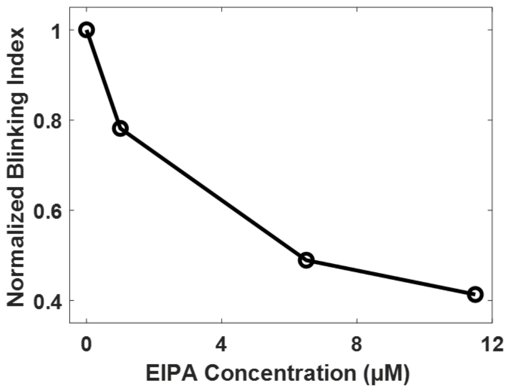


Fig. S7. Decreasing blinking index of bacteria with the increasing concentration of EIPA.

#### 2.8 Synchronized BL blinking and fluctuations in membrane potential

Fig. S8 shows the correlative BL (black) and FL (red) recordings of two representative bacteria labeled by TMRM, a membrane potential sensitive dye. First, synchronized drops were observed in both curves, indicating that the BL blinking was accompanied with the depolarizations in membrane potential. Second, the peaks in BL channel were significantly narrower than those in FL channel, demonstrating a better temporal resolution of using BL blinking to report the fluctuations in membrane potentials. For example, multiple continuous blinking events (marked by blue stars in Fig. S8a) can be clearly observed and distinguished in BL channel. However, the FL channel only displays a broad peak. In addition, many blinking events with smaller amplitudes (marked by blue stars in Fig. S8b) can only be detected in BL channel. The comparisons suggested that BL emission could be a promising readout signal to reveal the membrane potential dynamics of single bacteria with improved temporal resolution and sensitivity.

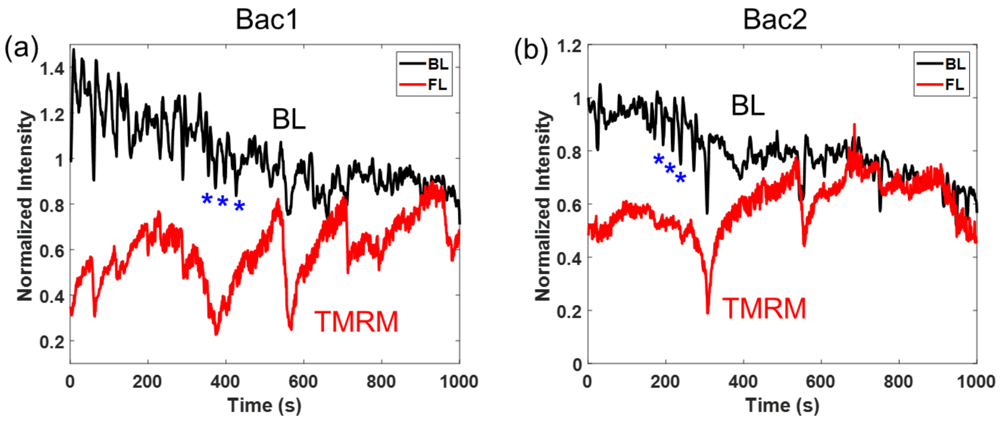


Fig. S8. Correlative BL (black curves) and FL (red curves) recordings of two representative bacteria labeled by TMRM (a, b), a membrane potential sensitive dye.

#### 2.9 Occasionally synchronized BL blinking and fluctuations in intracellular pH

We have introduced a ratiometric pH-sensitive fluorophore, RatioWorks™ PH165, NHS ester to investigate bacterial $pH_i$. While the BL trajectory exhibited frequent and obvious blinking features, the correlatively recorded $pH_i$ remained stable. It was reasonable because buffering capacity of neutrophilic bacteria was previously reported to be 10~100 nmol protons per pH unit per mg cell protein under normal physiological conditions[2]. The dry weight of a single bacterium measured was about 100 fg[3]. We assume that the protein content accounts for 30% of cell dry weight. As the result, the number of protons to alter 1 pH unit can be calculated as follows:

$$10 \sim 100\ \text{nmol/mg protein} \times 30\ \text{fg protein} \times 6.02 \times 10^{23}$$

In order to alter 1 pH unit in a single bacterium, $1.8\times10^{5\sim6}$ protons were needed.

In some rare cases, $pH_i$ curve was dropped simultaneously with BL signal. The falling extent of $pH_i$ was rather small, less than 0.1 pH unit. It suggested that the proton was moved into the cytoplasm when blinking events happened, but $pH_i$ was nearly constant because of the greater buffer capacity of cytoplasmic substances.

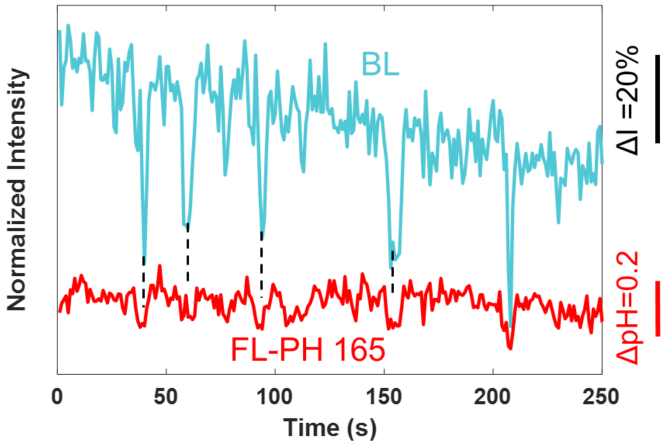


Fig. S9. Correlative BL (blue curves) and FL (red curves) recordings of a single bacterium labeled by RatioWorks™ PH165, NHS ester, a ratiometric fluorescent probe for intracellular pH.

#### 2.10 Distribution histograms of blinking depths at five temperatures

The conformation change of $Na^+/H^+$ antiporter was driven by thermal fluctuation. As a result, the activation energy barrier of this process can be calculated by analyzing blinking depth at different temperatures. We adopted temperature control system[1] to generate heat, and statistically analyzed the blinking depth of hundreds of single bacteria at five different temperatures. The distribution histograms of blinking depths (Fig. S10) demonstrated the increasing inhibition ratio of higher temperatures.

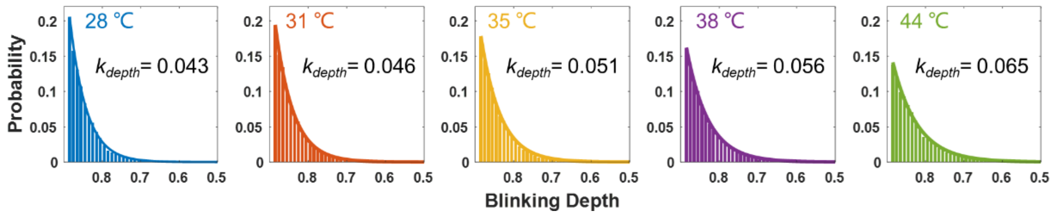


**Fig. S10.** The distribution histograms of blinking depths at five temperatures.

#### 2.11 The dependance of $\tau_{on}$ and $k_{depth}$

$\tau_{on}$ and $k_{depth}$ were used to describe the duration times of ON events and the inhibition depth of BL blinking, respectively. We calculated the correlative $\tau_{on}$ and $k_{depth}$ at five solutions of pH. The linear dependence of $\tau_{on}$ and $k_{depth}$ was shown in Fig. S11. The result showed that longer $\tau_{on}$ led to larger inhibition ratio ($k_{depth}$) with the increasing pH values.

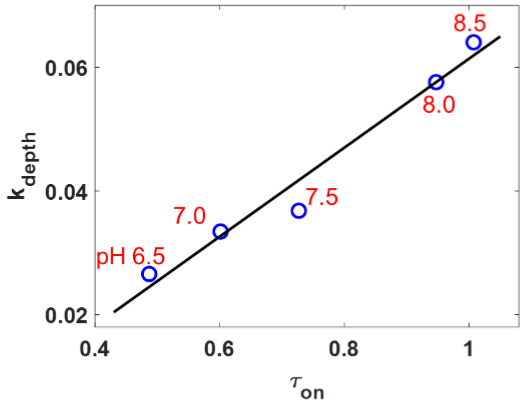


**Fig. S11.** The dependence of $\tau_{on}$ and $k_{depth}$ at different solutions of pH.

#### 2.12 Measurements of intracellular pH

We used a commercial ratiometric pH-sensitive fluorophore, RatioWorks™ PH165, NHS ester, to measure bacterial $pH_i$. Its emission wavelength does not coincide with bacterial BL emission. Moreover, it shows pH-dependent fluorescence spectra. This probe was popularly used to quantify $pH_i$ in the range from 4 to 9 with dual excitation/emission at 497/594 nm and 578/654 nm, respectively.

An intracellular pH calibration curve was made by adding 10 μM nigericin into the medium to equalize cytoplasmic and extracellular pHs. The dependence of intracellular pH and ratios of FL intensity ($Em_{594}/Em_{654}$) was shown in Fig. S12a. Similarly, bacteria in the culture mediums of pH4.0-9.0 without nigericin were also measured. According to the relationship shown in Fig. S12a, intracellular pH ($pH_i$) as a function of extracellular pH ($pH_e$) was obtained for native bacteria. It was clear that $pH_i$ was generally closer to neutral condition when the medium was too acidic ($pH_e < 6$) or too basic ($pH_e > 8$), indicating the buffering capacity of the cytoplasm.

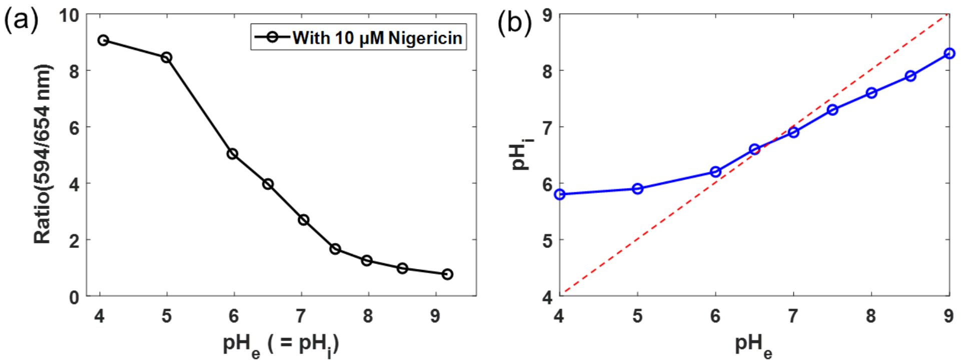


Fig. S12. (a) The dependence of FL ratiometric ($Em_{594}/Em_{654}$) on $pH_e$ in the presence of 10 μM nigericin. Because nigericin was able to equalize cytoplasmic and extracellular pHs ($pH_e = pH_i$), this curve acted as the calibration curve to determine $pH_i$ in the absence of nigericin (b).

#### 2.13 Effects of $H^+$-ATPase inhibitors

The plasma membrane $H^+$-ATPase plays an essential role in establishing and maintaining the membrane potential by harvesting the energy from ATP hydrolysis to pump protons out of the cells. In order to examine the roles of $H^+$-ATPase in BL blinking, the influences of two $H^+$-ATPase inhibitors, oligomycin, and N, N'-dicyclohexylcarbodiimide (DCCD), were investigated. As shown in Fig. S13, with the increasing concentrations of these two inhibitors, the blinking indexes accordingly decreased.

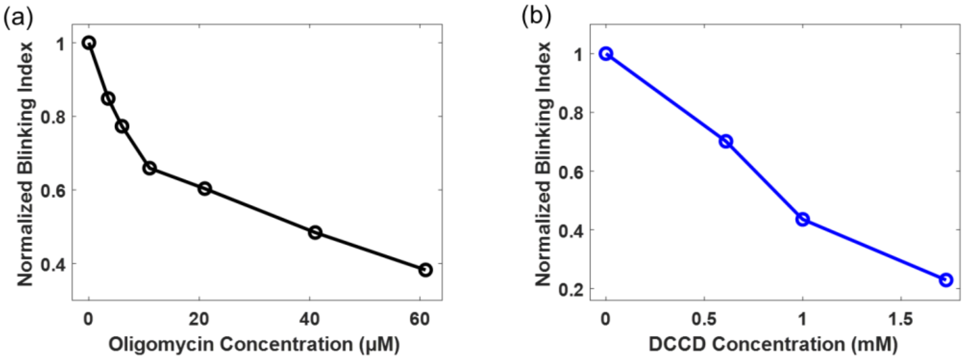


Fig. S13. Decreasing blinking index of bacteria with the increasing concentrations of oligomycin (a) and DCCD (b), respectively.

#### 2.14 BL blinking in wild type *V. fischeri*

pH-dependent BL blinking was also observed in another kind of bioluminescent bacteria *V. fischeri.* Blinking index was increased by 17 times with a pH rise from 7.0 to 8.0. These results demonstrated the generality of bacterial BL blinking.

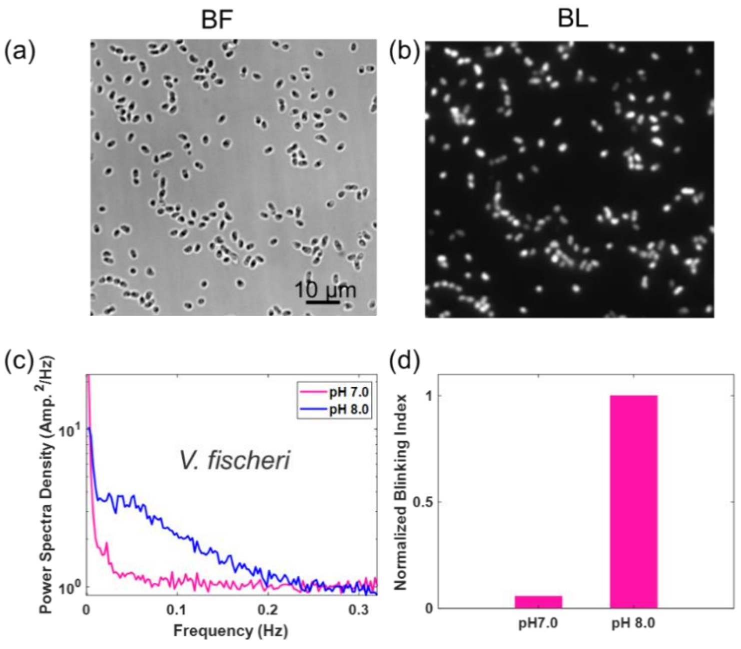

Fig. S14. Correlative BF (a) and BL (b) images of *V. fischeri* adhered on the coverslip. PSD curves (c) and the corresponding blinking index (d) were extracted from time-lapsed BL images of *V. fischeri* in the solution of pH 7.0 (pink) and 8.0 (blue), respectively.

#### Supplementary Tables

In order to examine the existence of NhaA gene in the native *P. phosphoreum* strain we used in the study, gene sequencing analysis was performed by Sangon Biotech Co., Ltd. (Shanghai, China). The result showed that *P. phosphoreum* indeed contained the NhaA gene.

**Table S1. Gene and amino acid sequences of NhaA in *P. phosphoreum***

**Gene sequence:**

ATGACCGATGCTATTCGTAAGTTTCTTAAATTAGAATCAGCTGGCGGTATTATTCTTA
TAATCGCAGCATTAATTGCAATGGTTATTGCTAATTCACCATTAGCATCTGTATATAC
TGACACTTTGCATAGCTATATTGCAGGATTATCAGTAGCGCATTGGATCAATGATGG
CTTAATGGCGATTTTCTTTTTCTTAATTGGATTAGAAGTTAAGCGTGAACTTATTGAG
GGTGCACTTAATACTAAGGAAAAAGCAATATTTCCAGCTATTGCTGCTGTCGGCGGT
ATGATTGCGCCAGCATTAATTTATACCGCATTTAATTACGGCGATCCGATGGCGGTT
AAAGGCTGGGCAATTCCTGCGGCGACAGATATTGCTTTTGCACTTGGTGTGATGGC
ATTACTTGGTAACCGTGTTCCTGTGAGCTTAAAAGTTTTCTTATTAGCCTTGGCTATT
ATTGATGACTTAGGTGTGATTGTAATCATTGCTTTATTTTATAGCTCTGACTTATCGA
CTATCGCGCTGACAGTGGCATTTGTCGCTACCGCCACCTTAATCATTATGAATCTAA
AGAATGTCACTAGCATTCCATTATATTTGATTGTCGGTGCTATTTTATGGTTCAGCGT
GCTTCAATCTGGTGTGCATGCAACATTAGCTGGTGTGGTGCTAGGTTTTGCCGTGC
CATTAGCCAGTAAAGATGGTCGTACAGATACTCATTCGCCGCTTAAAACGATTGAAC
ATGCGCTACATCCATATGTTGCCTTCTTAATCTTGCCGTTATTTGCGTTTGCGAATGC
TGGTATTTCACTAACGGGCGTATCACTGGCGAGTTTATCAGCCATGTTACCGGTTGG
TATTGCAGCGGGACTATTTATTGGTAAGCCAGTCGGTATATTTACTGCATGTTATATT
GCAGTTAAAACAGGCGTTGCAAAATTGCCTGATGGGATTAACTTCAAGCATATTTTT
GCGGTTTCTGTGCTGTGTGGTATCGGCTTTACGATGTCTATCTTCATCTCATCACTG
GCGTTTGTTGGTGCAGATGAGAGTTTTGCAACTTATTCACGCCTTGGAATCTTATTT
GGTTCGACGGTTGCCGCGATTGTTGGTTACATAATGTTGAGTAAAACACTACCTAAA
TCAGAGGTGTAA

**Amino acid sequence:**

MTDAIRKFLKLESAGGIILIIAALIAMVIANSPLASVYTDTLHSYIAGLSVAHWINDGLMAIFF
FLIGLEVKRELIEGALNTKEKAIFPAIAAVGGMIAPALIYTAFNYGDPMAVKGWAIPAATDI
AFALGVMALLGNRVPVSLKVFLLALAIIDDLGVIVIIALFYSSDLSTIALTVAFVATATLIIMNL
KNVTSIPLYLIVGAILWFSVLQSGVHATLAGVVLGFAVPLASKDGRTDTHSPLKTIEHALH
PYVAFLILPLFAFANAGISLTGVSLASLSAMLPVGIAAGLFIGKPVGIFTACYIAVKTGVAKL
PDGINFKHIFAVSVLCGIGFTMSIFISSLAFVGADESFATYSRLGILFGSTVAAIVGYIMLSK
TLPKSEV

#### Supplementary Movies

**Movie S1.** BL blinking of *P. phosphoreum* in the culture medium of pH8.0 was recorded with a 2 s exposure time. The play rate of this movie is 100x fast forward.

**Movie S2.** BL blinking of *P. phosphoreum* in the culture medium of pH 6.5 and pH 8.5 was recorded with an exposure time of 1 s. The play rate of this movie is 100x fast forward.

**Movie S3.** BL blinking of *V. fischeri* in the culture medium of pH8.0 was recorded with a 2 s exposure time. The play rate of this movie is 100x fast forward.
